## Supplemental Materials for "Placental Network Differences Among Obstetric Syndromes Identified With An Integrated Multiomics Approach"

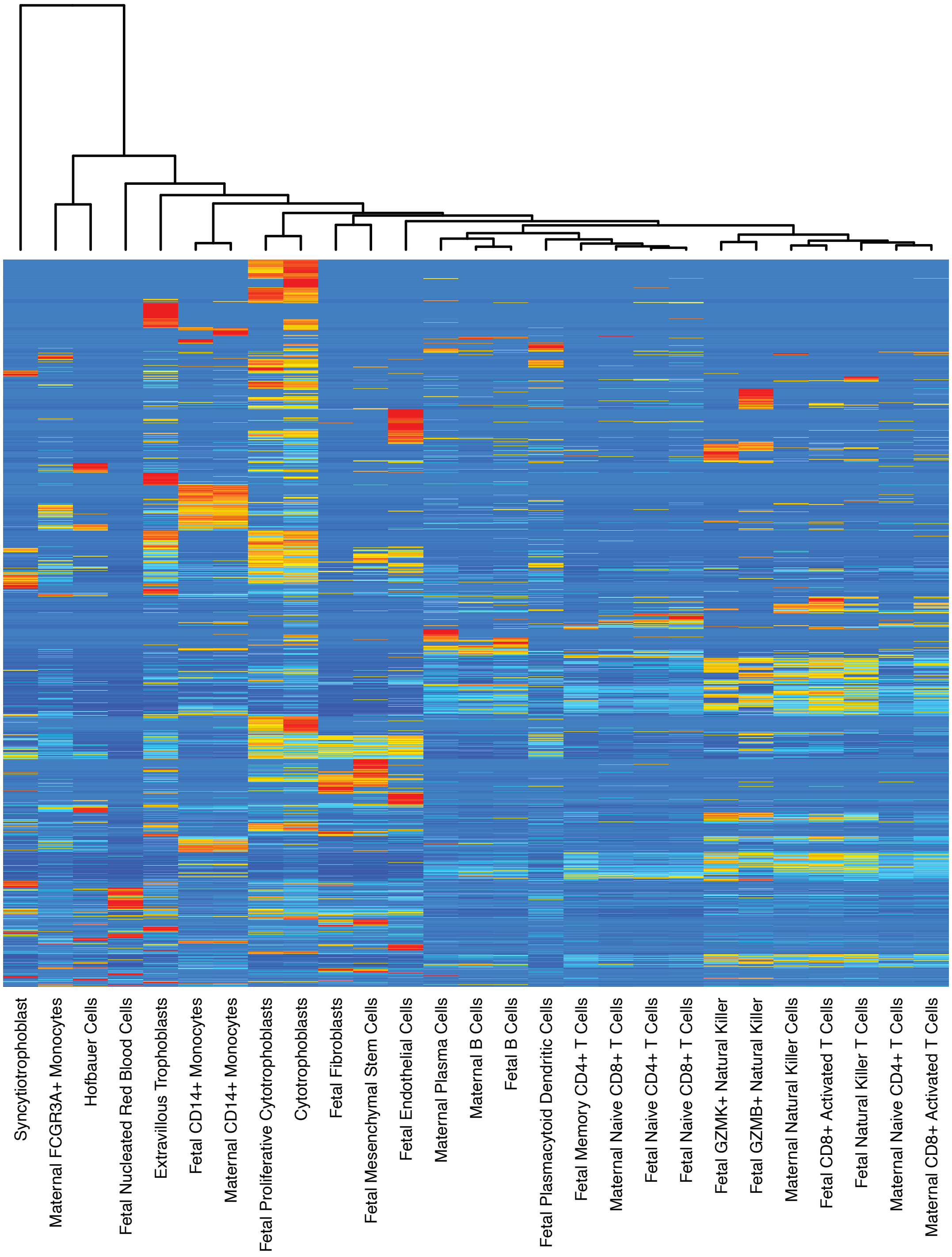

**Supplemental Figure 1. Gene expression across placental cell types.** Cell type cluster map of cell types (x-axis) and gene expression (y-axis). The color indicates the level of gene expression within a given cell type with red indicating high level of expression and blue indicating low level of expression. Hierarchical clustering was performed to group both the cell types for similarity of gene expression. This was generated from the cell-type signature matrix was generated using a single-cell RNA sequencing reference sample[1] normalized to counts per million (CPM), batch corrected using “S-mode” and permuted 100 times to assess statistical significance. This was generated by CIBERSORTx.

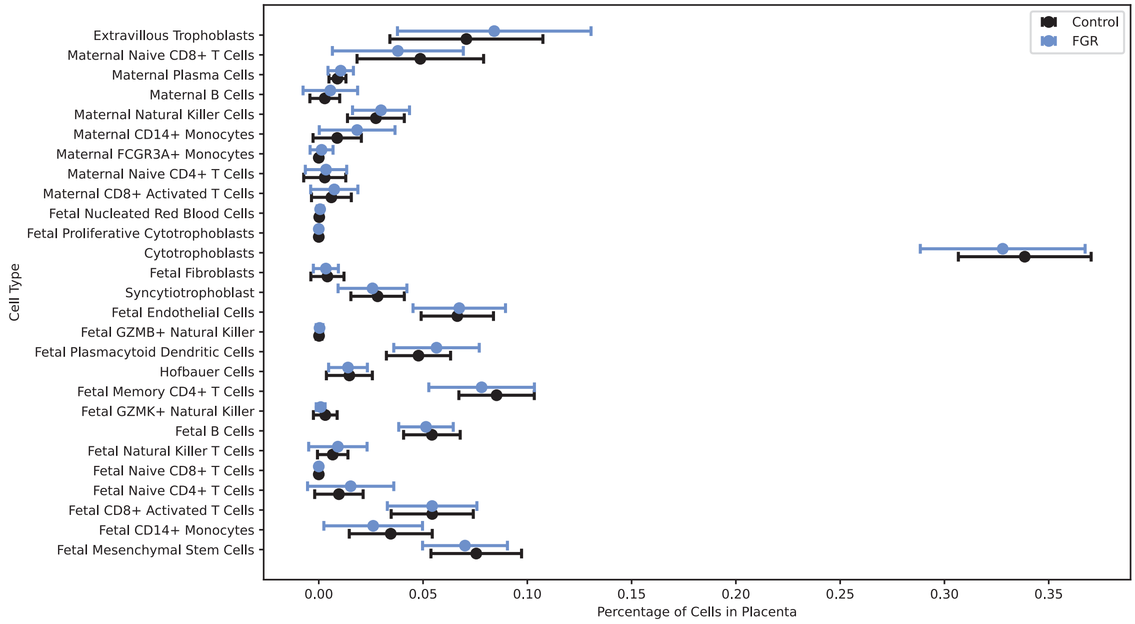

**Supplemental Figure 2. No difference in cell type composition in FGR placentas.** Dot plot with error bars of the mean cell type percentage plus standard deviation. The cell type percentages displayed were not significantly different (p>0.05) in any FGR (blue) compared to control (black) placentas. The p-values were calculated by Kolmogorov-Smirnov tests followed by Benjamini-Hochberg multiple hypothesis corrections.

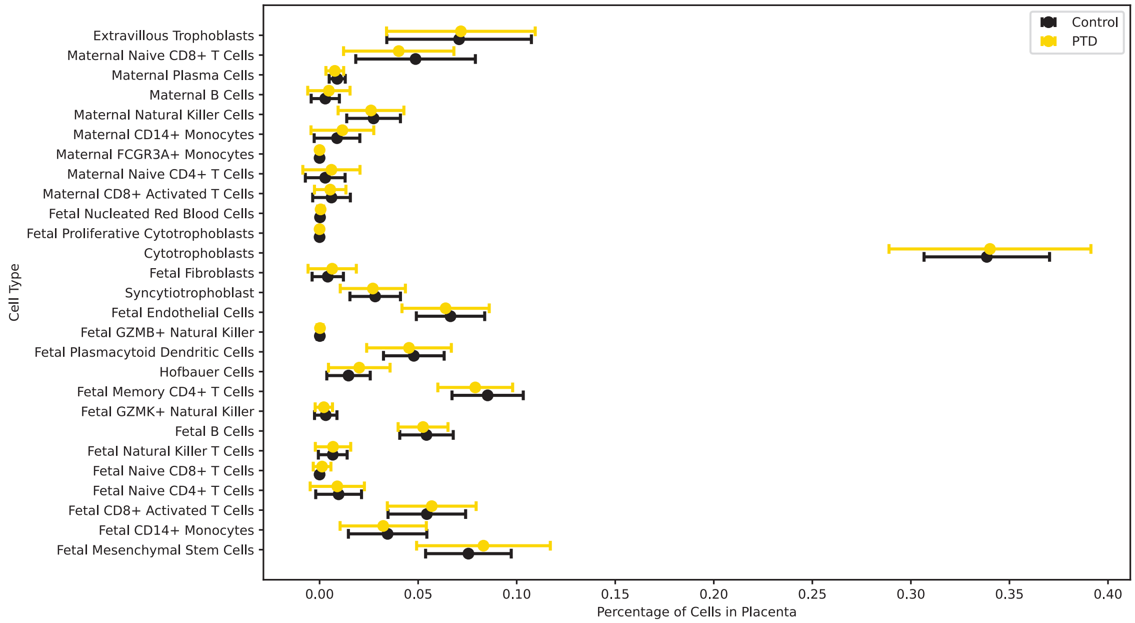

**Supplemental Figure 3. No difference in cell type composition in PTD placentas.** Dot plot with error bars of the mean cell type percentage plus standard deviation. The cell type percentages displayed were not significantly different (p>0.05) in any PTD (gold) compared to control (black) placentas. The p-values were calculated by Kolmogorov-Smirnov tests followed by Benjamini-Hochberg multiple hypothesis corrections.

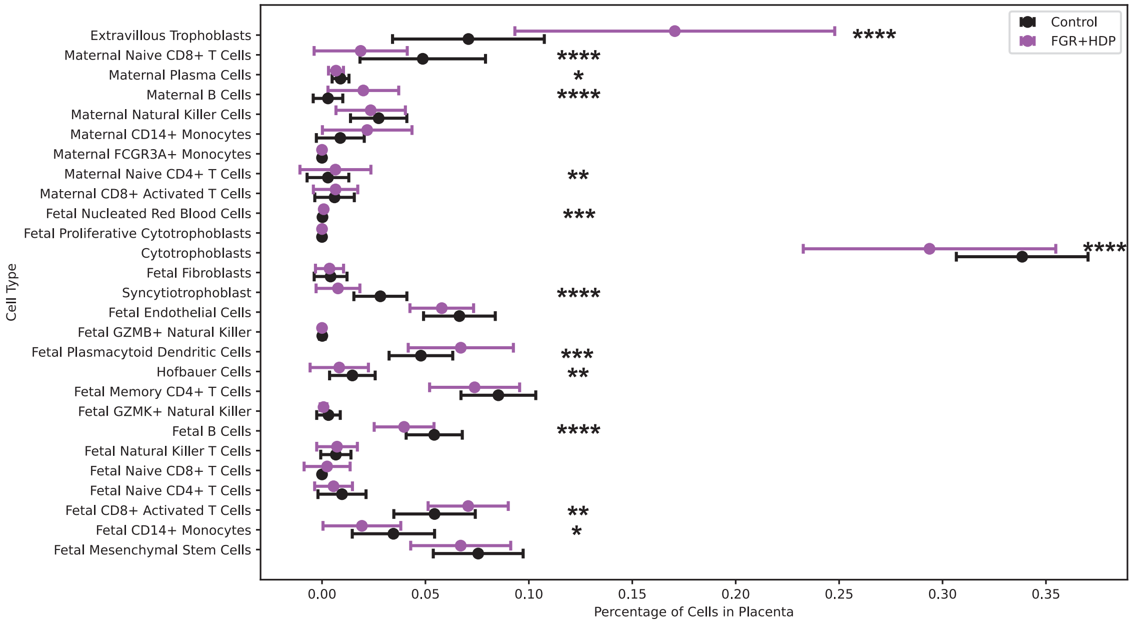

**Supplemental Figure 4. Cell type composition differences in FGR+HDP placentas.** Dot plot with error bars of the mean cell type percentage plus standard deviation. For a subset of cell types, the cell type percentages displayed were significantly different in FGR+HDP (purple) compared to control (black) placentas. The p-values were calculated by Kolmogorov-Smirnov tests followed by Benjamini-Hochberg multiple hypothesis corrections. *p<0.05, **p<0.01, ***p<0.001, ****p<0.0001.

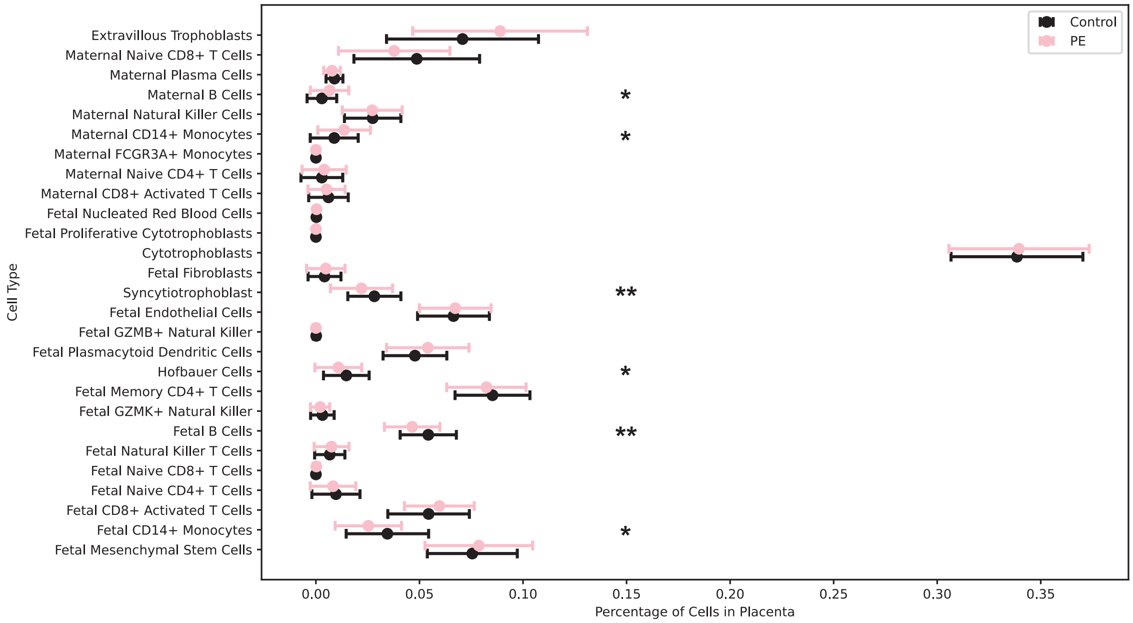

**Supplemental Figure 5. Cell type composition differences in PE placentas.** Dot plot with error bars of the mean cell type percentage plus standard deviation. For a subset of cell types, the cell type percentages displayed were significantly different in PE (pink) compared to control (black) placentas. The p-values were calculated by Kolmogorov-Smirnov tests followed by Benjamini-Hochberg multiple hypothesis corrections. *p<0.05, **p<0.01.

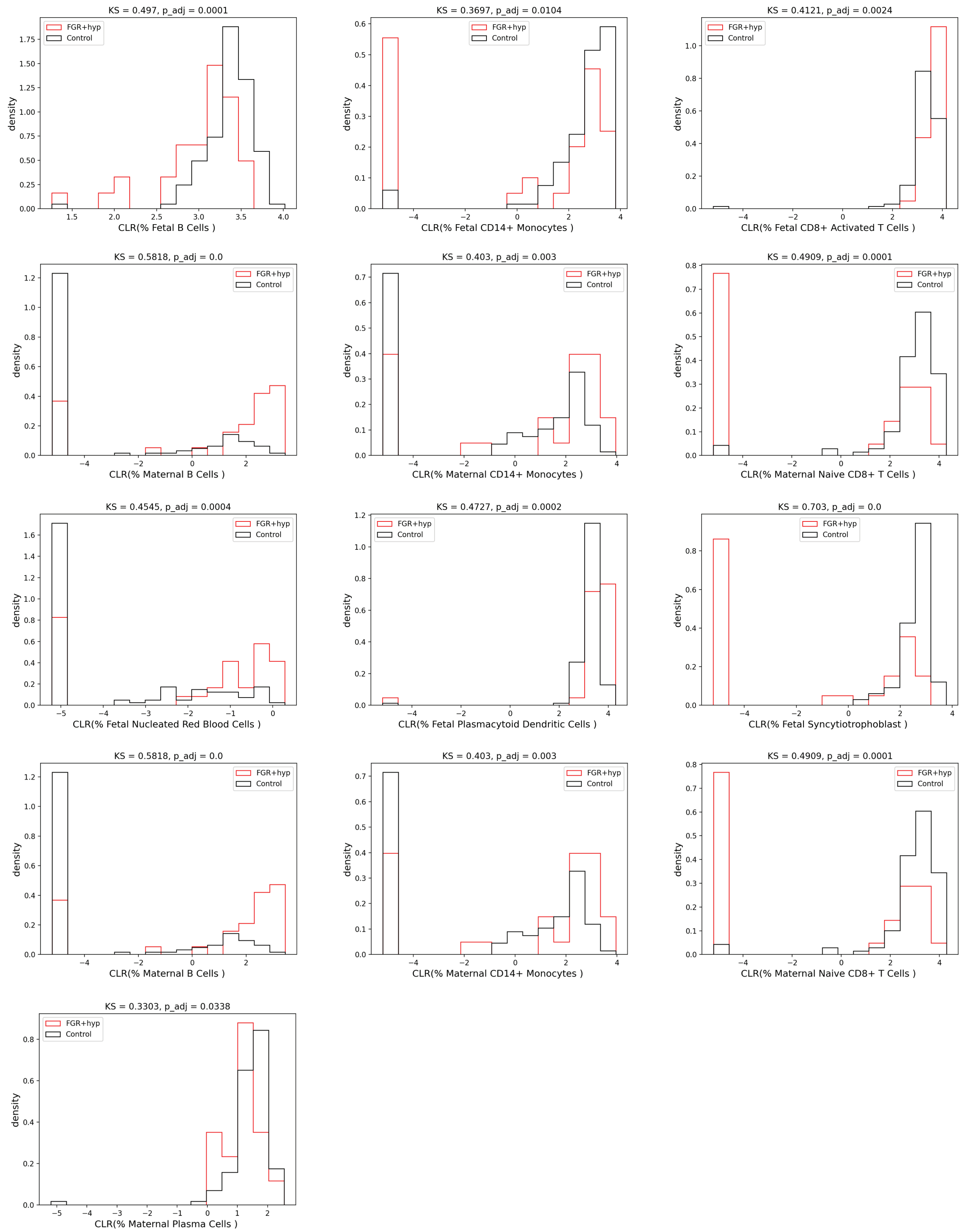

**Supplemental Figure 6. CLR-transformed cell type distributions in FGR+HDP placentas**. Histograms of clr-transformed cell type composition in FGR+HDP (red) and control (black) placentas. The p-values were calculated by Kolmogorov-Smirnov tests followed by Benjamini-Hochberg multiple hypothesis corrections.

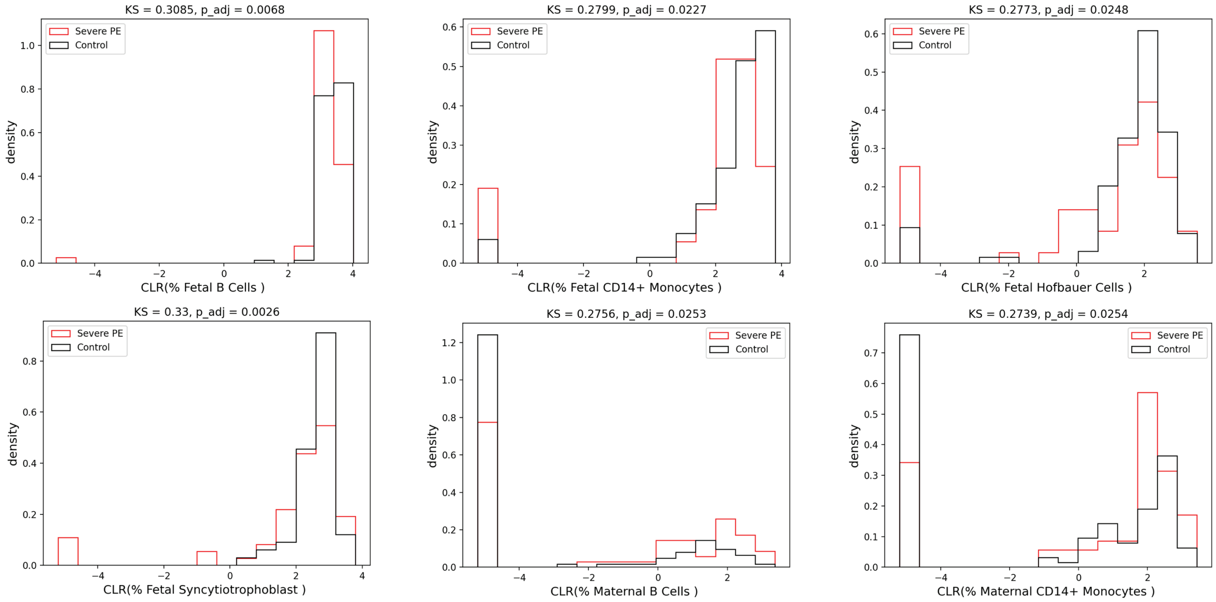

**Supplemental Figure 7. CLR-transformed cell type distributions in PE placentas.** Histograms of clr-transformed cell type composition in PE (red) and control (black) placentas. The p-values were calculated by Kolmogorov-Smirnov tests followed by Benjamini-Hochberg multiple hypothesis corrections.

### Female Fetuses

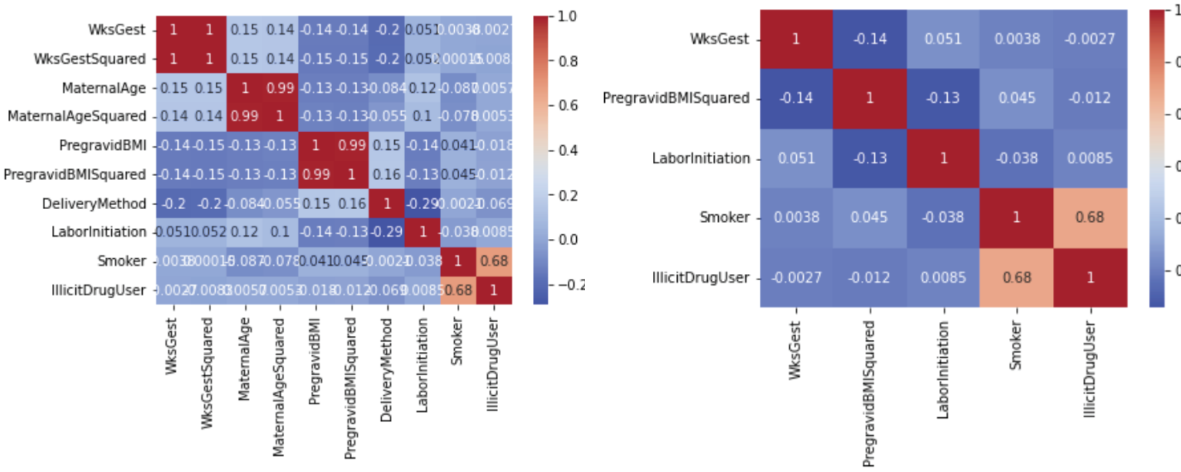

### Male Fetuses

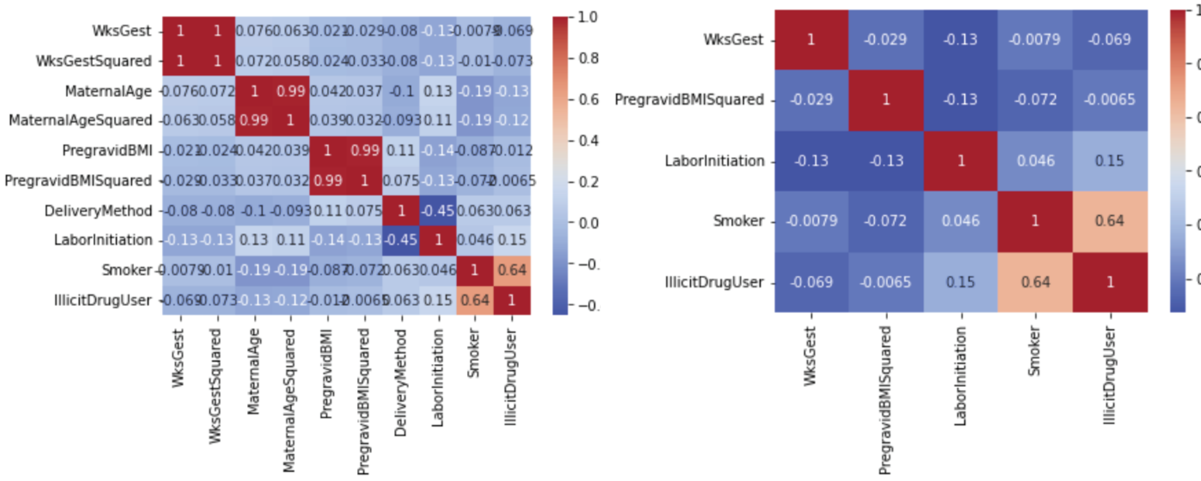

**Supplemental Figure 8. Correlation matrices of common confounders.** Correlation matrices to evaluate common confounders in female (top) and male (bottom) fetuses. Two matrices are displayed for each sex: one with all the confounding variables considered (left) and one containing only the variables used in the generalized linear models (GLM; right). The color indicates the strength of the correlation with dark blue indicating no correlation (0) and dark red indicating perfect correlation (1).

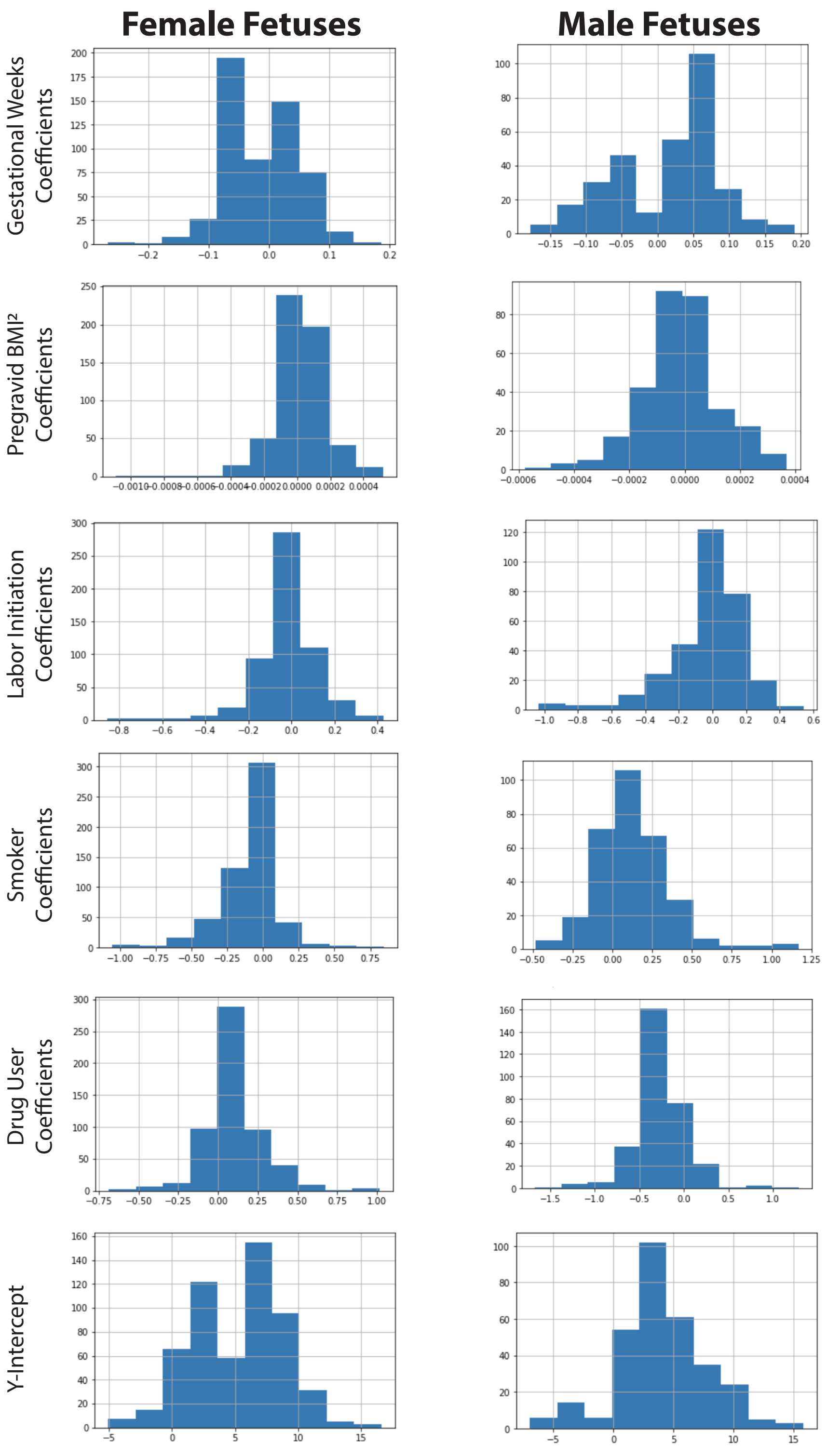

**Supplemental Figure 9. Effect sizes are small for cofactors.** Histograms of the β-coefficients for each variable in the GLMs for female (left) and male (right) fetuses. The β-coefficients were limited to those in GLMs whose gestational weeks at delivery was significantly regulated (FDR<0.05) following Benjamini-Hochberg multiple hypothesis adjustment

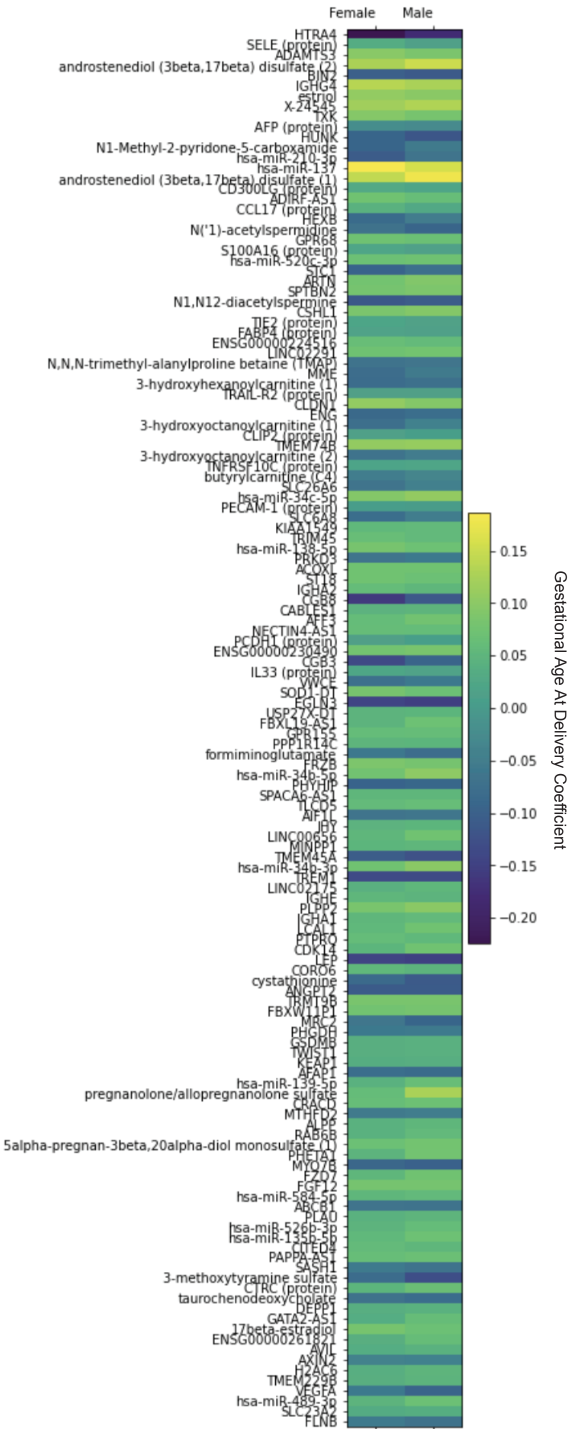

**Supplemental Figure 10. Gestational weeks at delivery effect size is comparable between fetal sexes.** Heatmap of the β-coefficient of the gestational weeks at delivery for the GLMs of analytes that are significantly regulated (FDR<0.05) by this variable in both fetal sexes following Benjamini-Hochberg multiple hypothesis adjustment. Purple indicates a low, negative β-coefficient and yellow indicates a high, positive β-coefficient.

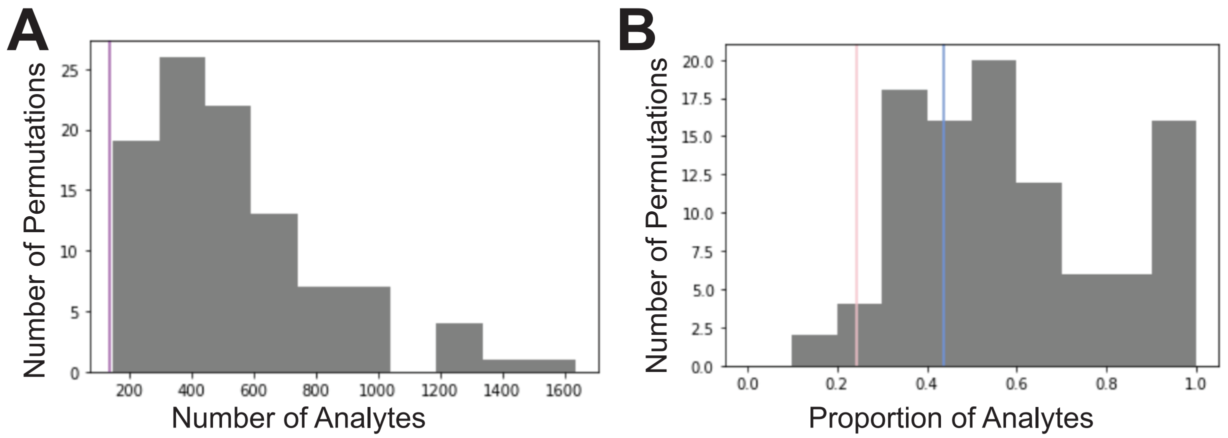

**Supplemental Figure 11. Random simulation of overlap of analytes significantly regulated by gestational age at delivery between two groups**. A hundred iterations were performed in which each sample was assigned to one of two groups, GLMs were fit for each analyte in each group, and the overlap in the analytes significantly regulated (FDR<0.05) by gestational weeks at delivery in both random groups following Benjamini-Hochberg multiple hypothesis correction was assessed. (**A**) Histogram (bin size=100) of the number of significant analytes in common between the two groups across all permutations. The purple vertical line (y=136) is what is observed when comparing female and male fetuses. As this falls below the range of anything observed by random chance this means FDR<0.01 for this observation. (**B**) Histogram (bin size=0.1) of the proportion of analytes significantly regulated by gestational weeks at delivery in a random group that was observed to be in common with the second random group across all permutations. The vertical pink (y=0.244) and blue (y=0.439) lines represent the proportion of significant analytes observed in female and male fetuses respectively. Where these fall within the range of random permutations indicates a FDR<0.02 and FDR=0.14 for the proportion of significant analytes in female fetuses that also are significantly regulated in male fetuses and vice-versa.

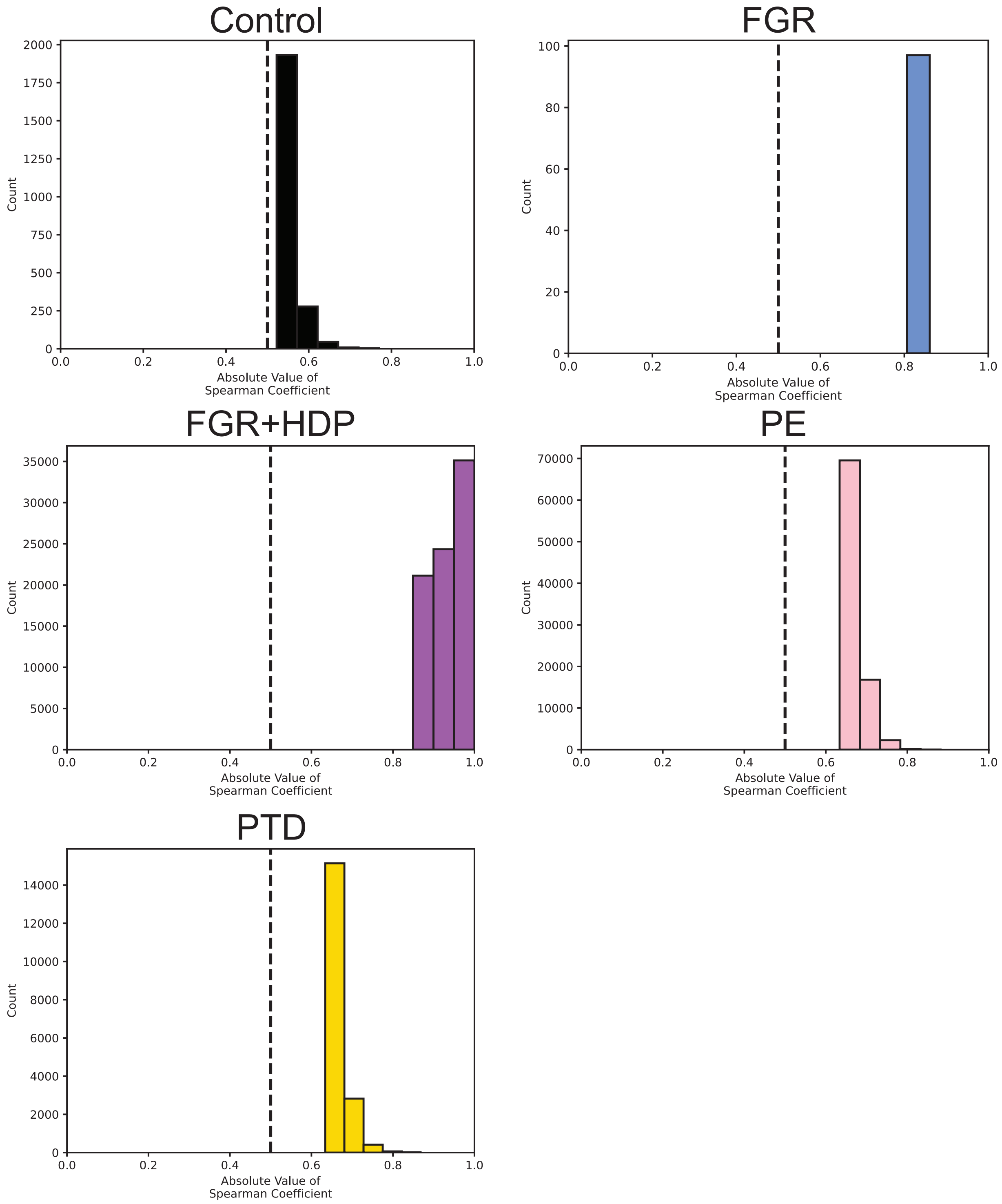

**Supplemental Figure 12. Effect size of significant interomics correlations is large across all obstetric conditions.** Histograms of the absolute value of the spearman coefficients of significant ($p<0.05$) interomics correlations following Bonferroni correction. The vertical dotted line ($y=0.5$) is the boundary for large effect size indicating that anything to the right of the line has a large effect size. This was reported for control (black), FGR (blue), FGR+HDP (purple), PE (pink), and PTD (gold) interomics networks.

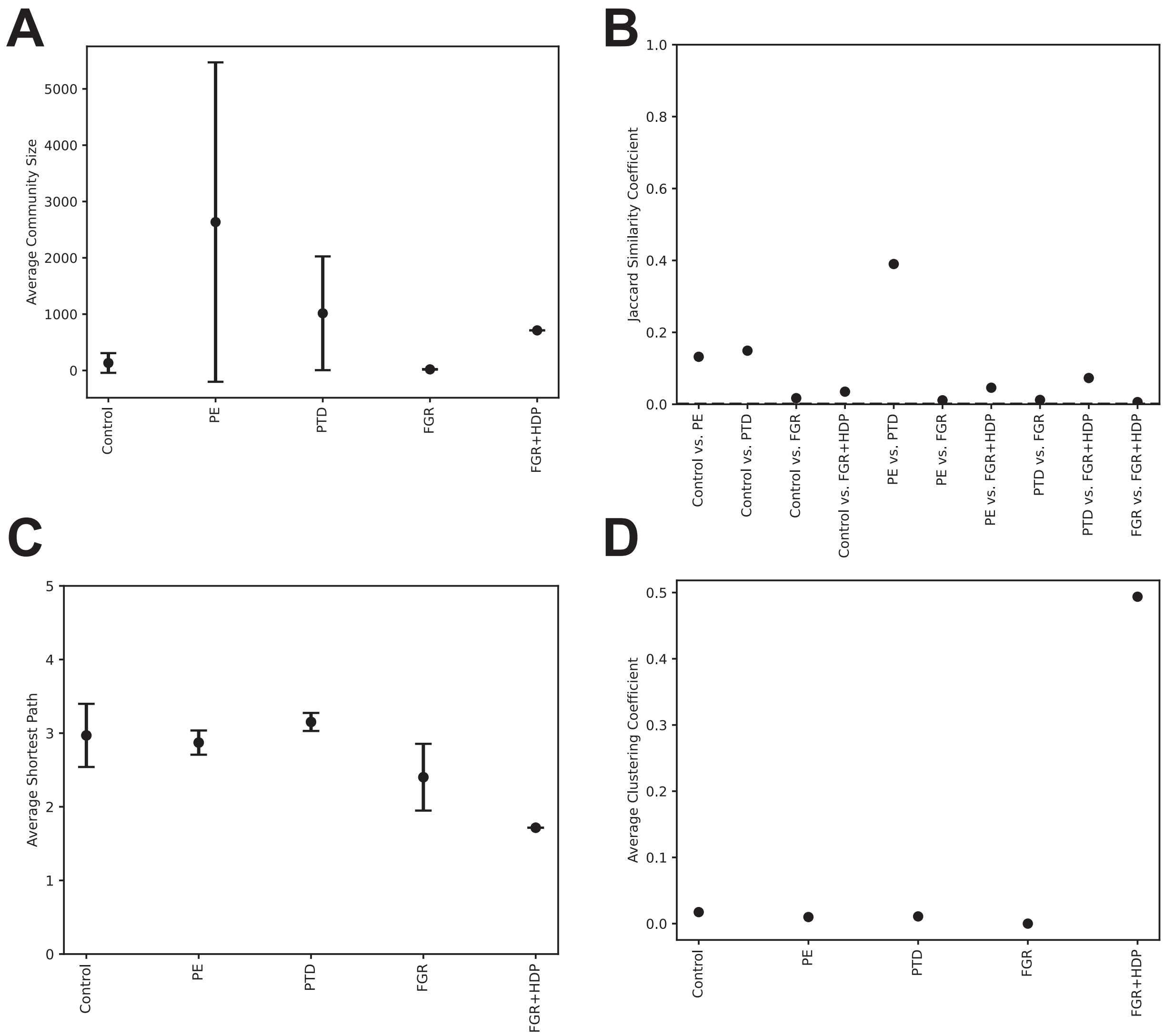

**Supplemental Figure 13. FGR+HDP had the densest and least structured network**. Evaluations of the interomics network structure and composition across obstetric conditions. (**A**) Dot plot with error bars of mean and standard deviation of the average community size. (**B**) Dot plots of the Jaccard Similarity Coefficient between all possible pairwise combinations of conditions. (**C**) Dot plot with error bars of the mean and standard deviation of the average shortest path within a community. (**D**) Dot plot of the average clustering coefficient.

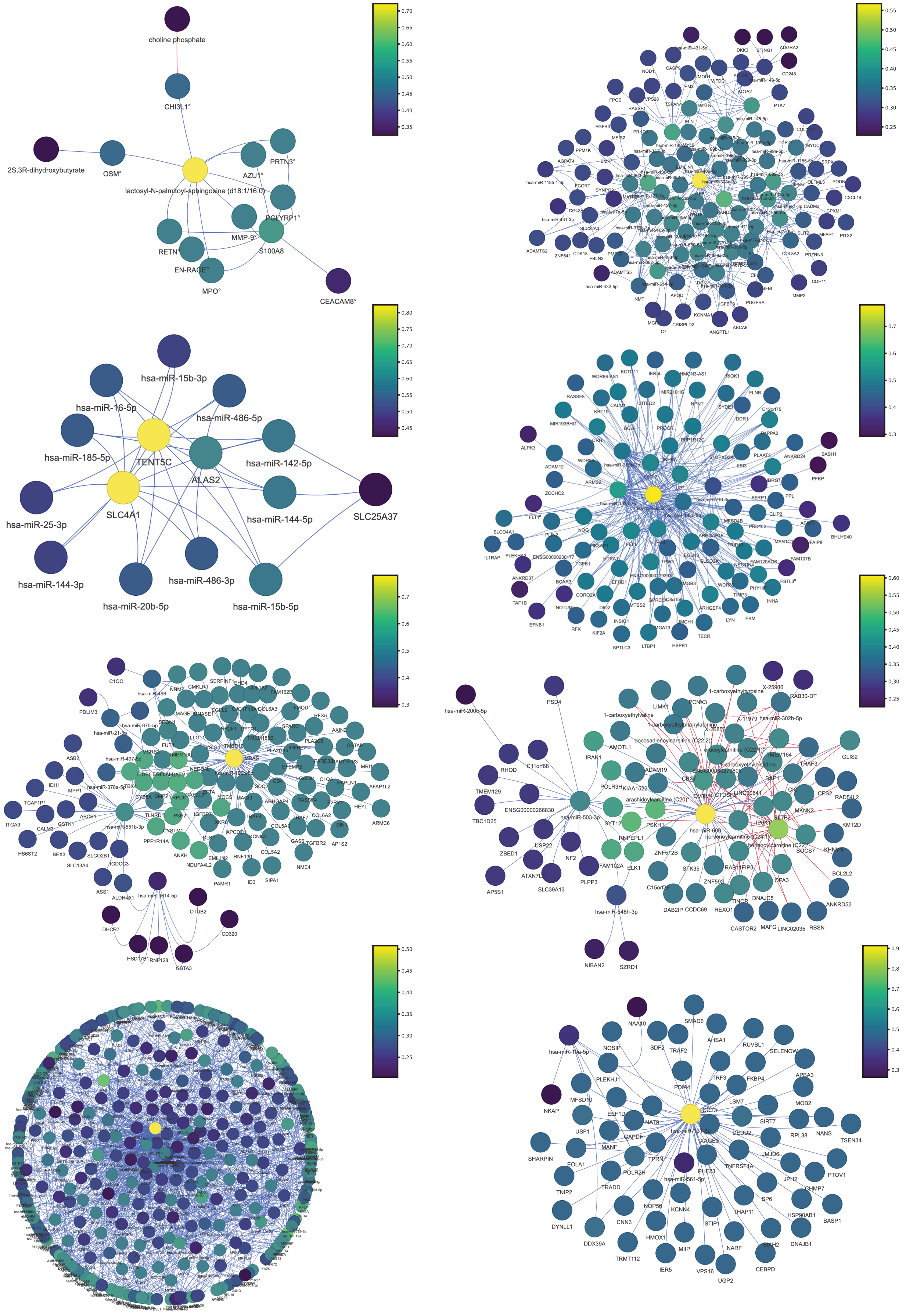
choline phosphate
CHI3L1°
2S,3R-dihydroxybutyrate
OSM°
lactosyl-N-palmitoyl-sphingosine (d18:1/16:0)
AZU1°
PRTN3°
PGLYRP1°
MMP-9°
S100A8
RETN°
EN-RAGE°
MPO°
CEACAM8°
0.70
0.65
0.60
0.55
0.50
0.45
0.40
0.35
hsa-miR-15b-3p
hsa-miR-16-5p
hsa-miR-486-5p
hsa-miR-185-5p
TENT5C
hsa-miR-142-5p
ALAS2
SLC4A1
hsa-miR-25-3p
hsa-miR-144-5p
SLC25A37
hsa-miR-144-3p
hsa-miR-486-3p
hsa-miR-20b-5p
hsa-miR-15b-5p
0.80
0.75
0.70
0.65
0.60
0.55
0.50
0.45

**Supplemental Figure 14. Control interomics communities**. All control interomics communities that have ten or more nodes. Analyte names are listed below each node. A ° was added to the end of protein names (e.g. FLT1°) to distinguish proteins from mRNAs. The edge color indicates the directionality of the correlation with blue representing a positive correlation and red representing a negative correlation The node color indicates the closeness centrality with yellow indicating the most connected node and purple indicating the least connected node. A color bar is to the right of each community and was normalized to a scale of 0 to 1 for each community. The individual html files for each of these communities is available on the paper's GitHub repository.

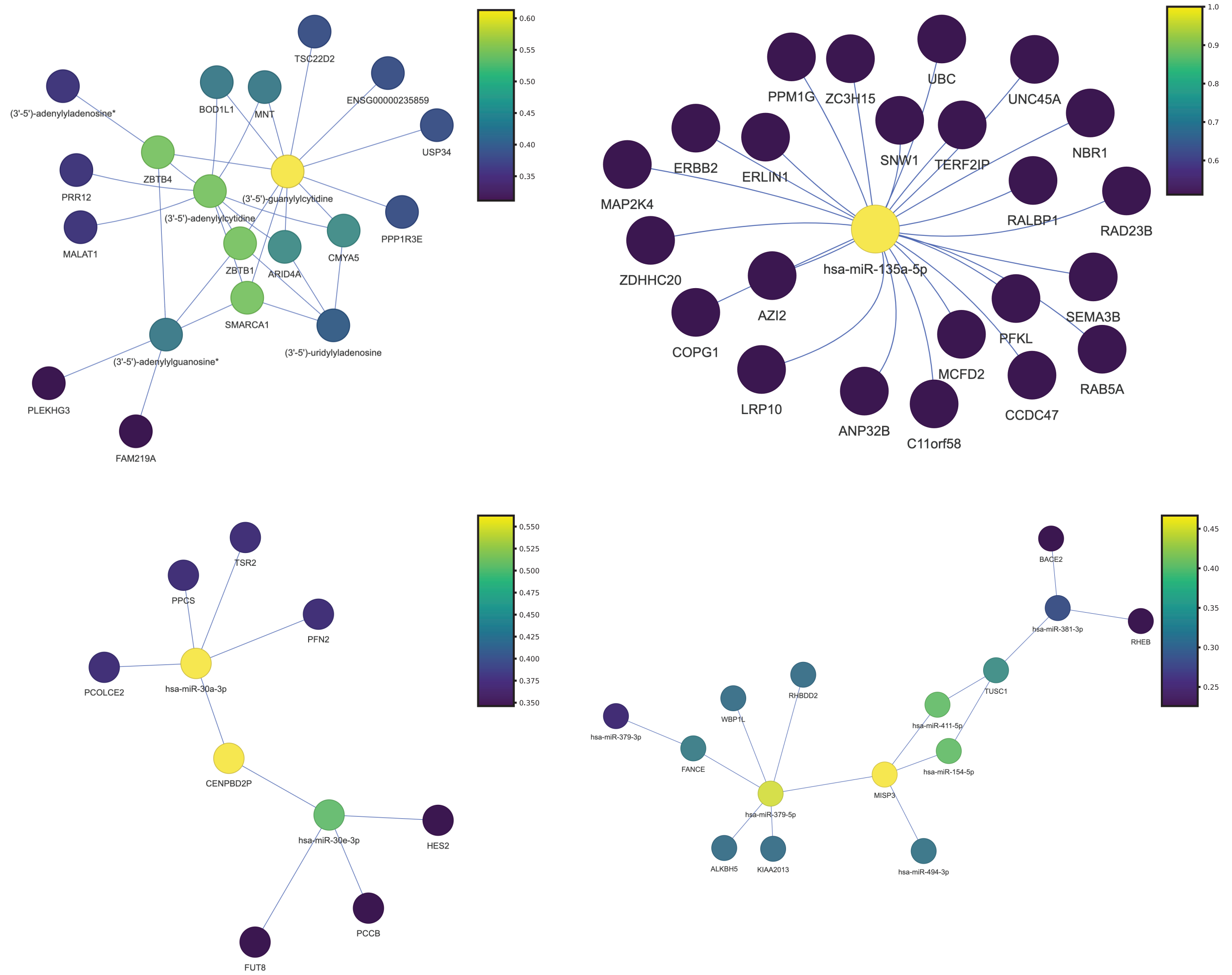

**Supplemental Figure 15. FGR interomics communities.** All FGR interomics communities that have ten or more nodes. Analyte names are listed below each node. A ° was added to the end of protein names (e.g. FLT1°) to distinguish proteins from mRNAs. The edge color indicates the directionality of the correlation with blue representing a positive correlation and red representing a negative correlation The node color indicates the closeness centrality with yellow indicating the most connected node and purple indicating the least connected node. A color bar is to the right of each community and was normalized to a scale of 0 to 1 for each community. The individual html files for each of these communities is available on the paper's GitHub repository.

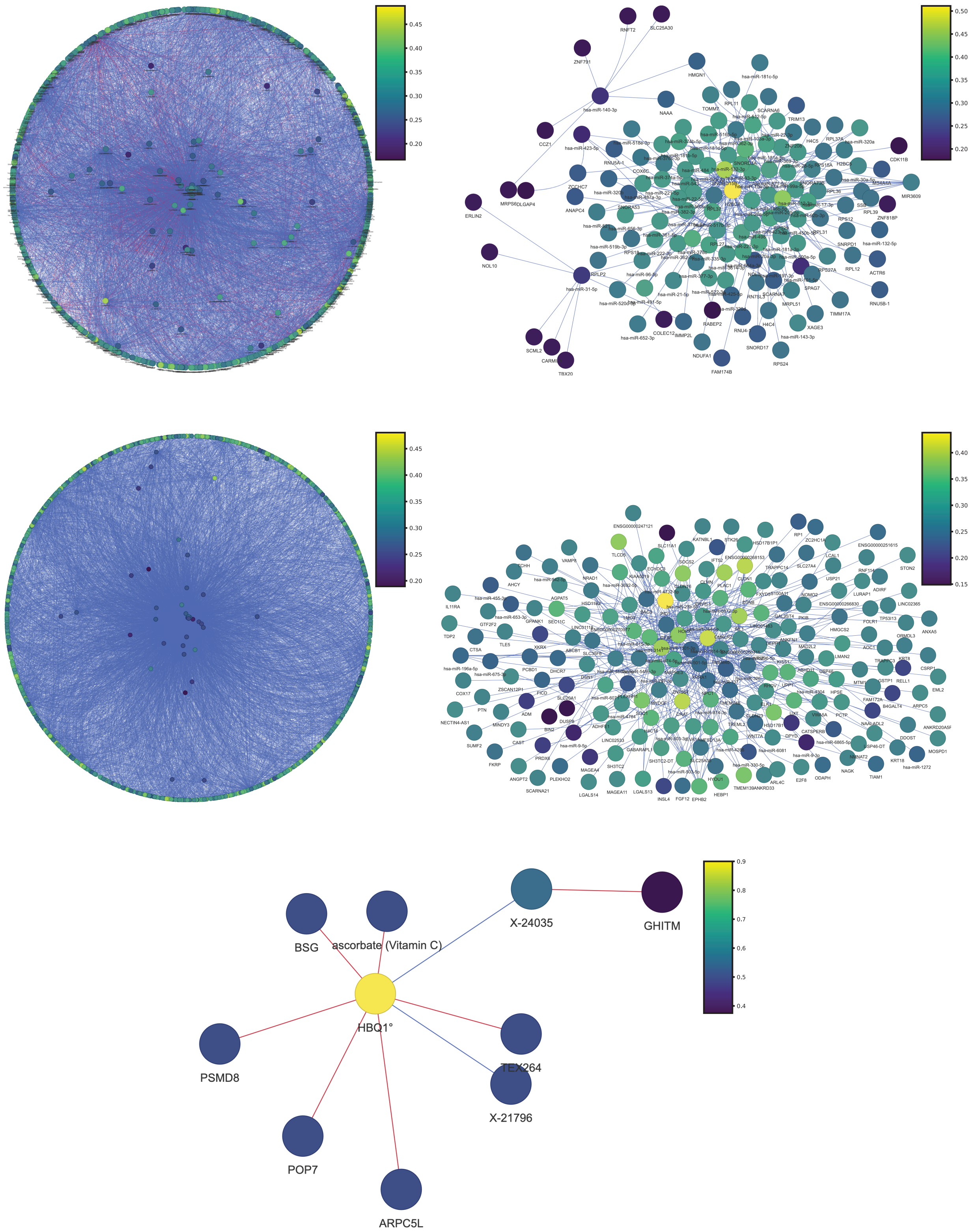
BSG
ascorbate (Vitamin C)
X-24035
GHITM
HBQ1°
PSMD8
TEX264
X-21796
POP7
ARPC5L
0.9
0.8
0.7
0.6
0.5
0.4

**Supplemental Figure 16. PTD interomics communities**. All PTD interomics communities that have ten or more nodes. Analyte names are listed below each node. A ° was added to the end of protein names (e.g. FLT1°) to distinguish proteins from mRNAs. The edge color indicates the directionality of the correlation with blue representing a positive correlation and red representing a negative correlation The node color indicates the closeness centrality with yellow indicating the most connected node and purple indicating the least connected node. A color bar is to the right of each community and was normalized to a scale of 0 to 1 for each community. The individual html files for each of these communities is available on the paper's GitHub repository.

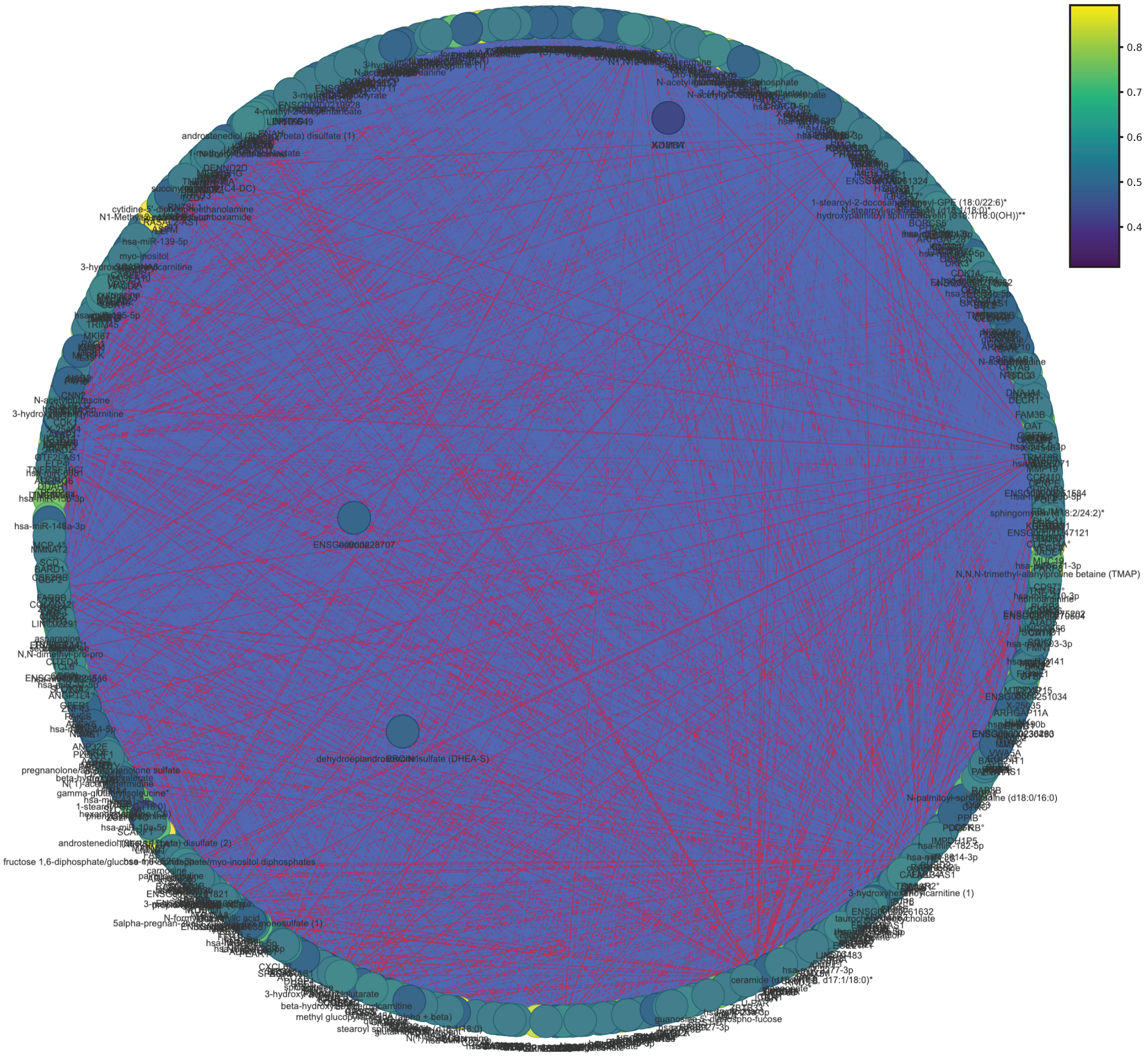

**Supplemental Figure 17. FGR+HDP interomics community**. The only FGR+HDP interomics community that has ten or more nodes. Analyte names are listed below each node. A ° was added to the end of protein names (e.g. FLT1°) to distinguish proteins from mRNAs. The edge color indicates the directionality of the correlation with blue representing a positive correlation and red representing a negative correlation The node color indicates the closeness centrality with yellow indicating the most connected node and purple indicating the least connected node. The color bar was normalized to a scale of 0 to 1 for each community. The individual html files for this community is available on the paper's GitHub repository.

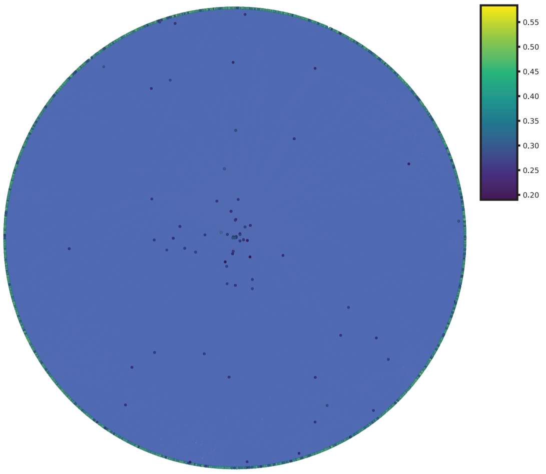

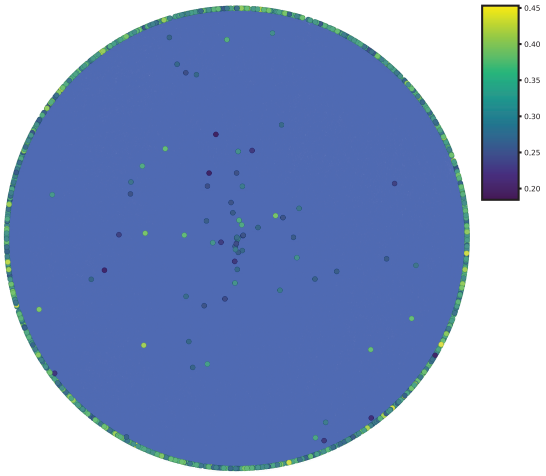

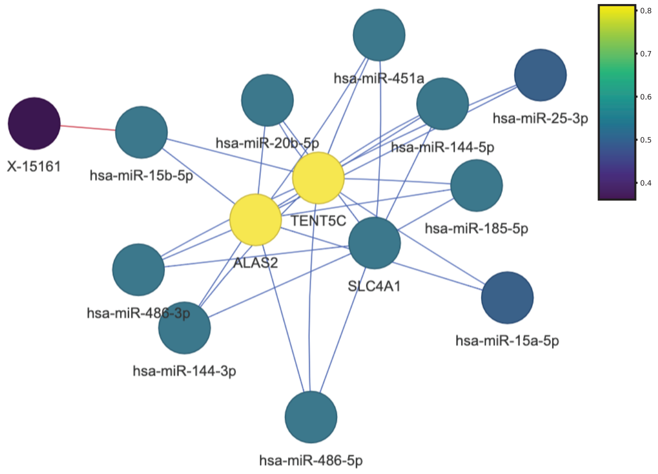

**Supplemental Figure 18. PE interomics communities.** All PE interomics communities that have ten or more nodes. Analyte names are listed below each node. A ° was added to the end of protein names (e.g. FLT1°) to distinguish proteins from mRNAs. The edge color indicates the directionality of the correlation with blue representing a positive correlation and red representing a negative correlation The node color indicates the closeness centrality with yellow indicating the most connected node and purple indicating the least connected node. A color bar is to the right of each community and was normalized to a scale of 0 to 1 for each community. The individual html files for each of these communities is available on the paper's GitHub repository.

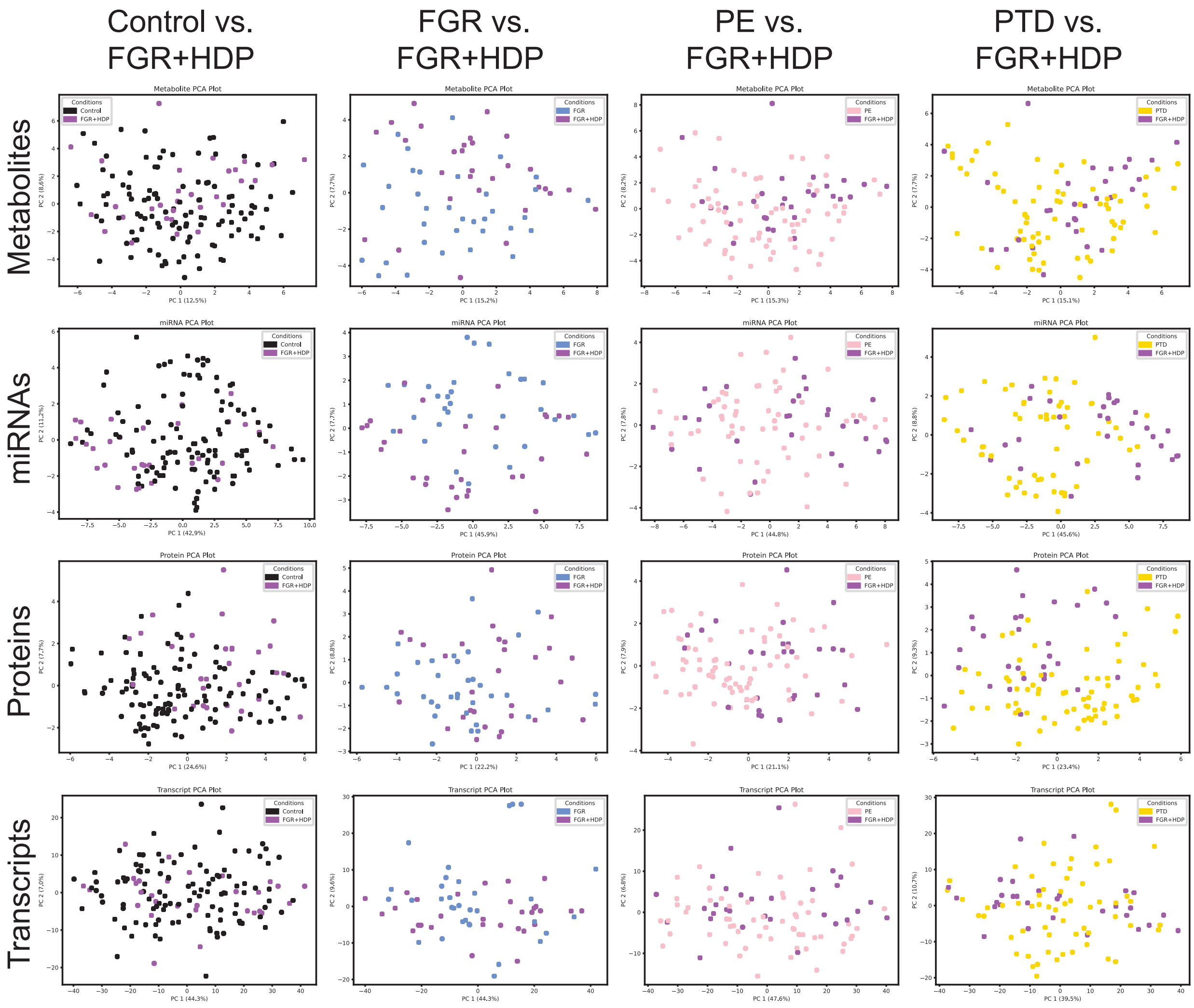

**Supplemental Figure 19. Cannot distinguish between FGR+HDP and other obstetric conditions using all measured analytes**. Principal Component Analyses (PCA) first two dimensions were plotted using all metabolites (first row), miRNAs (second row), proteins (third row), and mRNA transcripts (fourth row). This was done for FGR+HDP (purple) alongside each of the other obstetric conditions: Control (black; column one), FGR (blue; column two), PE (pink, column three), and PTD (gold, column four). Each dot represents a placenta and the color of the dot indicates the condition to which it belongs.

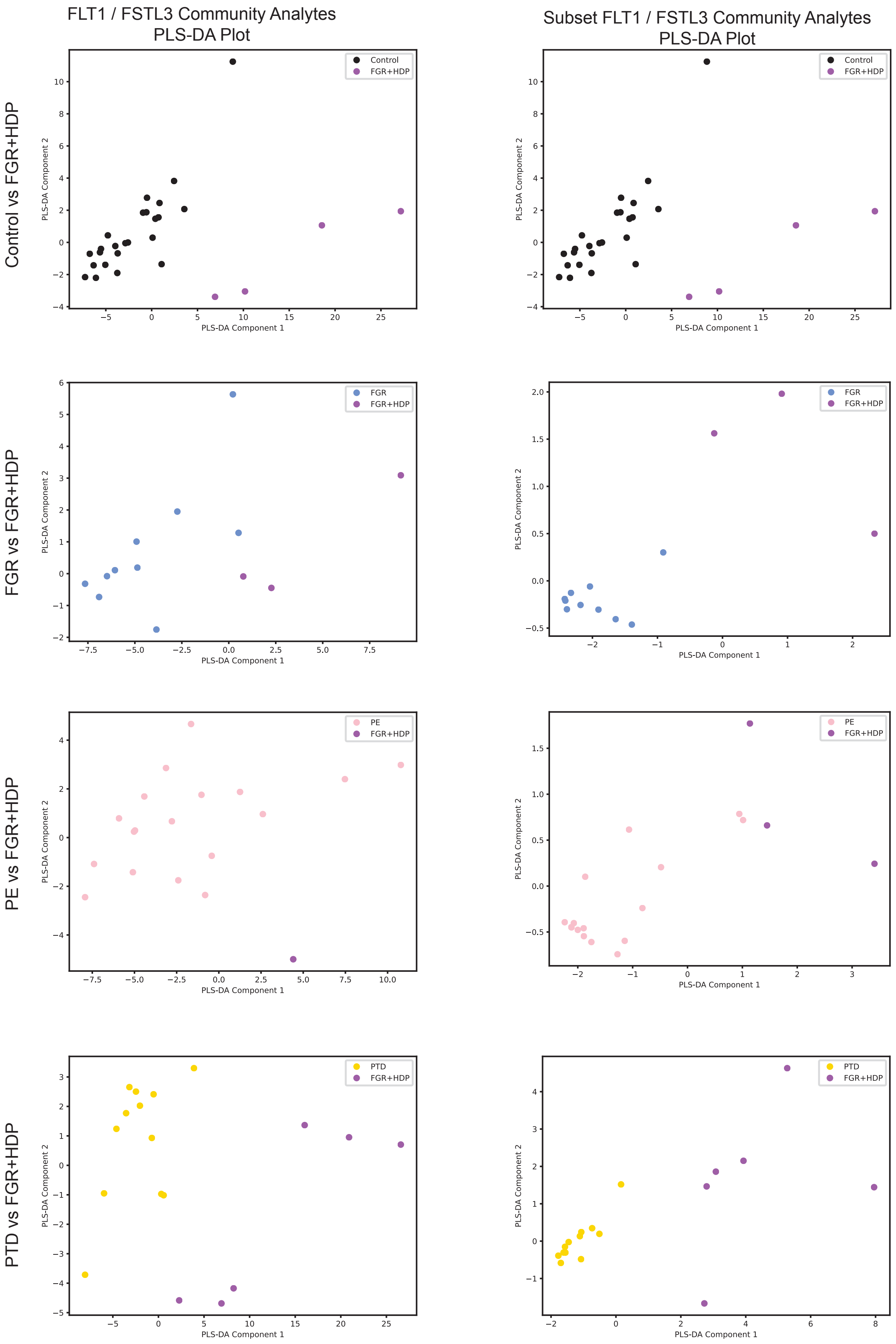
FLT1 / FSTL3 Community Analytes
PLS-DA Plot
Subset FLT1 / FSTL3 Community Analytes
PLS-DA Plot
Control vs FGR+HDP
FGR vs FGR+HDP
PE vs FGR+HDP
PTD vs FGR+HDP
Control
FGR
PE
PTD
FGR+HDP
PLS-DA Component 1
PLS-DA Component 2

**Supplemental Figure 20. FGR+HDP signature distinguishes it from placentas with overlapping clinical features.** Partial Least Square-Discriminant Analysis (PLS-DA) of the FGR+HDP (purple) placentas plotted alongside the control (black; top panels), FGR (blue; middle top panels), PE (pink; middle bottom panels), and PTD (gold; bottom panels) for both all 100 analytes in FLT1/FSTL3 community (left) and the twelve mRNA transcripts in the final FGR+HDP biosignature (right). Each dot represents a placenta with the color of the dot indicating the placenta's condition. The twelve analytes in the final FGR+HDP biosignature were selected for having the most distinctive differential expression across all pairwise comparisons of conditions as revealed by hierarchical clustering (Figure 4B).

| Feature | Definition |
| --- | --- |
| **Demographic** | |
| Maternal Age (Years) | Maternal age at start of pregnancy (years); continuous variable |
| Pregravid BMI (kg/m2) | Pregravid Body Mass Index (BMI; kg/m2); continuous variable |
| Race | Reported race: Asian, Black, Native American, White, Multiracial, or not reported (missing); categorical variable |
| Ethnicity, Hispanic | Reported ethnicity: Non-Hispanic or Unknown (0) or Hispanic (1); binary variable |
| **Social History** | |
| Smoking | Smoking status: not reported as smoking during pregnancy (0), or reported smoking during pregnancy (1) |
| Illicit Drug Use | Illicit drug status: no reported illicit drug use (0), or reported illicit drug use defined as cocaine use, substance abuse, marijuana use, maternal narcotic addiction at delivery, or drug withdrawal syndrome in newborn (1); binary variable |
| **Pregnancy Characteristics** | |
| Condition | Obstetric condition associated with the pregnancy: Control Term (delivery $\geq$37 weeks gestation, $\geq$10th percentile birthweight by gestational age, no pregnancy complications), Fetal Growth Restriction (FGR; <3rd percentile birthweight by gestational age; no hypertensive disorders); Fetal Growth Restriction with Precnancy Related Hypertension (FGR+HDP; <3rd percentile birthweight by gestational age with hypertensive disorder); Severe Preeclampsia (PE; severe features defined according to the American College of Obstetricians and Gynecologists guidelines); Spontaneous Preterm Delivery (PTD; delivery <37 weeks gestation with spontaneous labor present); categorical variable |
| Parity | Number of times a person has given birth to a fetus older than 24 weeks of gestation prior to the current pregnancy: Nulliparity (0 births), Multiparous (1 - 4 births) or Grand Multiparous (5+ births); categorical variable |
| **Delivery Charecteristics** | |
| Delivery Method | Reports method of delivery as being vaginal (0) or caesarean section (C-Section; 1); binary variable |
| Labor Initiation | Reports if spontaneous laber was present (0) or absent (1) at delivery, those without spontaneous labor may have been induced or had no labor at delivery; binary variable |
| Birthweight, g | Birthweight recorded at delivery (g); continuous variable |
| Gestational Age at Delivery, Weeks | Gestational age at delivery (weeks); continuous variable |

**Supplemental Table 1. Feature definitions for demographic, social history, pregnancy, and delivery characteristics**.

| | p-value |
|---|---|
| | Female vs Male Fetus |
| **Demographics** | |
| Maternal Age, Years | p=0.15 |
| Pregravid BMI, kg/m2 | p<0.01 |
| Race | p<0.05 |
| Ethnicity | p=0.97 |
| **Social History** | |
| Smoking | p=0.09 |
| Illicit Drug Use | p=0.53 |
| **Pregnancy Characteristics** | |
| Condition | p<0.01 |
| Parity | NA |
| **Delivery Characteristics** | |
| Delivery Method | p=0.06 |
| Labor Initiation | p<0.01 |
| Birthweight, g | p=0.32 |
| Gestational Weeks at Delivery | p=0.53 |

**Supplemental Table 2. Demographic, social history, pregnancy, and delivery characteristics composition between female and male fetuses**. Chi square test and students two tailed t-test were performed for categorical and continuous characteristics respectively.

| Analyte | Number Observed | Number Passing Cutoffs |
|---|---|---|
| Metabolites | 1032 | 865 |
| miRNAs | 2414 | 448 |
| Proteins | 452 | 343 |
| Transcripts | 51,174 | 9582 |
| Total | 52,658 | 11,238 |

**Supplemental Table 3. Number of analytes observed versus passing cutoffs by data type**.

| Cell Type | KS Statistic | p-value | Group Mean (CLR) | Control Mean (CLR) | Group STD (CLR) | Control STD (CLR) | Group Mean (%) | Control Mean (%) | Group STD (%) | Control STD (%) | p-value Adjusted |
|---|---|---|---|---|---|---|---|---|---|---|---|
| Fetal Mesenchymal Stem Cells | 0.184 | 0.287 | 3.616 | 3.693 | 0.300 | 0.286 | 0.070 | 0.076 | 0.020 | 0.022 | 0.535 |
| Fetal CD14+ Monocytes | 0.242 | 0.075 | 0.907 | 2.482 | 3.294 | 1.667 | 0.026 | 0.035 | 0.024 | 0.020 | 0.225 |
| Fetal CD8+ Activated T Cells | 0.096 | 0.945 | 3.247 | 3.262 | 0.743 | 0.914 | 0.054 | 0.054 | 0.021 | 0.020 | 1.000 |
| Fetal Naive CD4+ T Cells | 0.145 | 0.572 | -0.486 | -0.746 | 3.571 | 3.388 | 0.015 | 0.010 | 0.021 | 0.012 | 0.824 |
| Fetal Naive CD8+ T Cells | 0.061 | 1.000 | -5.196 | -5.196 | 0.000 | 0.000 | 0.000 | 0.000 | 0.000 | 0.000 | 1.000 |
| Fetal Natural Killer T Cells | 0.145 | 0.572 | -1.271 | -0.822 | 3.510 | 3.205 | 0.009 | 0.007 | 0.014 | 0.007 | 0.824 |
| Fetal B Cells | 0.149 | 0.540 | 3.286 | 3.364 | 0.450 | 0.307 | 0.051 | 0.054 | 0.013 | 0.014 | 0.821 |
| Fetal GZMK+ Natural Killer | 0.192 | 0.244 | -3.769 | -2.858 | 2.397 | 3.105 | 0.001 | 0.003 | 0.002 | 0.006 | 0.499 |
| Fetal Memory CD4+ T Cells | 0.213 | 0.154 | 3.691 | 3.828 | 0.438 | 0.248 | 0.078 | 0.085 | 0.025 | 0.018 | 0.377 |
| Fetal Hofbauer Cells | 0.092 | 0.959 | 1.615 | 1.500 | 1.425 | 1.816 | 0.014 | 0.015 | 0.009 | 0.011 | 1.000 |
| Fetal Plasmacytoid Dendritic Cells | 0.283 | 0.022 | 3.359 | 3.159 | 0.454 | 0.864 | 0.056 | 0.048 | 0.020 | 0.015 | 0.097 |
| Fetal GZMB+ Natural Killer | 0.061 | 1.000 | -4.828 | -5.130 | 1.496 | 0.690 | 0.000 | 0.000 | 0.002 | 0.001 | 1.000 |
| Fetal Endothelial Cells | 0.190 | 0.258 | 3.551 | 3.570 | 0.395 | 0.269 | 0.067 | 0.066 | 0.022 | 0.017 | 0.506 |
| Fetal Syncytiotrophoblast | 0.248 | 0.063 | 2.264 | 2.623 | 1.450 | 0.563 | 0.026 | 0.028 | 0.016 | 0.013 | 0.199 |
| Fetal Fibroblasts | 0.157 | 0.476 | -2.136 | -1.794 | 3.029 | 3.107 | 0.003 | 0.004 | 0.006 | 0.008 | 0.734 |
| Fetal Cytotrophoblasts | 0.229 | 0.105 | 5.195 | 5.229 | 0.120 | 0.097 | 0.328 | 0.339 | 0.040 | 0.032 | 0.280 |
| Fetal Proliferative Cytotrophoblasts | 0.061 | 1.000 | -5.196 | -5.196 | 0.000 | 0.000 | 0.000 | 0.000 | 0.000 | 0.000 | 1.000 |
| Fetal Nucleated Red Blood Cells | 0.240 | 0.078 | -2.929 | -3.827 | 2.204 | 1.868 | 0.001 | 0.000 | 0.001 | 0.000 | 0.226 |
| Maternal CD8+ Activated T Cells | 0.110 | 0.862 | -2.254 | -2.083 | 3.625 | 3.438 | 0.007 | 0.006 | 0.011 | 0.010 | 1.000 |
| Maternal Naive CD4+ T Cells | 0.078 | 0.992 | -3.867 | -3.920 | 2.750 | 2.673 | 0.003 | 0.003 | 0.010 | 0.010 | 1.000 |
| Maternal FCGR3A+ Monocytes | 0.075 | 0.995 | -4.753 | -5.196 | 1.801 | 0.000 | 0.001 | 0.000 | 0.005 | 0.000 | 1.000 |
| Maternal CD14+ Monocytes | 0.249 | 0.061 | 0.302 | -1.271 | 3.399 | 3.538 | 0.018 | 0.009 | 0.018 | 0.012 | 0.198 |
| Maternal Natural Killer Cells | 0.166 | 0.407 | 2.499 | 2.366 | 1.411 | 1.443 | 0.030 | 0.027 | 0.014 | 0.014 | 0.665 |
| Maternal B Cells | 0.139 | 0.629 | -2.642 | -3.348 | 3.245 | 2.959 | 0.006 | 0.003 | 0.013 | 0.007 | 0.886 |
| Maternal Plasma Cells | 0.157 | 0.476 | 1.583 | 1.445 | 0.643 | 0.795 | 0.011 | 0.009 | 0.006 | 0.004 | 0.734 |
| Maternal Naive CD8+ T Cells | 0.212 | 0.158 | 1.350 | 2.871 | 3.383 | 1.558 | 0.038 | 0.049 | 0.031 | 0.030 | 0.380 |
| Fetal Extravillous Trophoblasts | 0.177 | 0.335 | 3.689 | 3.526 | 0.572 | 0.568 | 0.084 | 0.071 | 0.046 | 0.037 | 0.593 |

**Supplemental Table 4. FGR cell type composition.** The cell type composition of FGR compared to control placentas. The mean and standard deviation are reported for each group as clr-transformed and percentage cell composition. The p-values were calculated by Kolmogorov-Smirnov tests followed by Benjamini-Hochberg multiple hypothesis corrections.

| Cell Type | KS Statistic | p-value | Group Mean (CLR) | Control Mean (CLR) | Group STD (CLR) | Control STD (CLR) | Group Mean (%) | Control Mean (%) | Group STD (%) | Control STD (%) | p-value Adjusted |
|---|---|---|---|---|---|---|---|---|---|---|---|
| Fetal Mesenchymal Stem Cells | 0.177 | 0.153 | 3.762 | 3.693 | 0.360 | 0.286 | 0.083 | 0.076 | 0.034 | 0.022 | 0.377 |
| Fetal CD14+ Monocytes | 0.112 | 0.668 | 2.046 | 2.482 | 2.330 | 1.667 | 0.032 | 0.035 | 0.022 | 0.020 | 0.890 |
| Fetal CD8+ Activated T Cells | 0.133 | 0.452 | 3.227 | 3.262 | 1.206 | 0.914 | 0.057 | 0.054 | 0.023 | 0.020 | 0.729 |
| Fetal Naive CD4+ T Cells | 0.156 | 0.269 | -1.669 | -0.746 | 3.611 | 3.388 | 0.009 | 0.010 | 0.014 | 0.012 | 0.519 |
| Fetal Naive CD8+ T Cells | 0.115 | 0.636 | -4.502 | -5.196 | 2.094 | 0.000 | 0.001 | 0.000 | 0.004 | 0.000 | 0.886 |
| Fetal Natural Killer T Cells | 0.121 | 0.572 | -1.343 | -0.822 | 3.366 | 3.205 | 0.007 | 0.007 | 0.009 | 0.007 | 0.824 |
| Fetal B Cells | 0.109 | 0.700 | 3.333 | 3.364 | 0.294 | 0.307 | 0.052 | 0.054 | 0.013 | 0.014 | 0.921 |
| Fetal GZMK+ Natural Killer | 0.123 | 0.557 | -3.005 | -2.858 | 2.932 | 3.105 | 0.002 | 0.003 | 0.004 | 0.006 | 0.824 |
| Fetal Memory CD4+ T Cells | 0.150 | 0.312 | 3.741 | 3.828 | 0.298 | 0.248 | 0.079 | 0.085 | 0.019 | 0.018 | 0.562 |
| Fetal Hofbauer Cells | 0.212 | 0.052 | 1.811 | 1.500 | 1.947 | 1.816 | 0.020 | 0.015 | 0.016 | 0.011 | 0.186 |
| Fetal Plasmacytoid Dendritic Cells | 0.159 | 0.250 | 2.767 | 3.159 | 1.878 | 0.864 | 0.045 | 0.048 | 0.021 | 0.015 | 0.499 |
| Fetal GZMB+ Natural Killer | 0.077 | 0.958 | -5.000 | -5.130 | 1.076 | 0.690 | 0.000 | 0.000 | 0.001 | 0.001 | 1.000 |
| Fetal Endothelial Cells | 0.208 | 0.060 | 3.514 | 3.570 | 0.325 | 0.269 | 0.064 | 0.066 | 0.022 | 0.017 | 0.198 |
| Fetal Syncytiotrophoblast | 0.144 | 0.359 | 2.239 | 2.623 | 1.777 | 0.563 | 0.027 | 0.028 | 0.017 | 0.013 | 0.616 |
| Fetal Fibroblasts | 0.171 | 0.181 | -1.230 | -1.794 | 3.178 | 3.107 | 0.006 | 0.004 | 0.012 | 0.008 | 0.426 |
| Fetal Cytotrophoblasts | 0.141 | 0.385 | 5.224 | 5.229 | 0.181 | 0.097 | 0.340 | 0.339 | 0.051 | 0.032 | 0.648 |
| Fetal Proliferative Cytotrophoblasts | 0.094 | 0.848 | -5.196 | -5.196 | 0.000 | 0.000 | 0.000 | 0.000 | 0.000 | 0.000 | 1.000 |
| Fetal Nucleated Red Blood Cells | 0.098 | 0.806 | -3.685 | -3.827 | 2.105 | 1.868 | 0.001 | 0.000 | 0.002 | 0.000 | 1.000 |
| Maternal CD8+ Activated T Cells | 0.071 | 0.980 | -1.998 | -2.083 | 3.384 | 3.438 | 0.005 | 0.006 | 0.008 | 0.010 | 1.000 |
| Maternal Naive CD4+ T Cells | 0.114 | 0.652 | -3.521 | -3.920 | 3.129 | 2.673 | 0.006 | 0.003 | 0.015 | 0.010 | 0.890 |
| Maternal FCGR3A+ Monocytes | 0.094 | 0.848 | -5.196 | -5.196 | 0.000 | 0.000 | 0.000 | 0.000 | 0.000 | 0.000 | 1.000 |
| Maternal CD14+ Monocytes | 0.086 | 0.907 | -0.807 | -1.271 | 3.494 | 3.538 | 0.012 | 0.009 | 0.016 | 0.012 | 1.000 |
| Maternal Natural Killer Cells | 0.097 | 0.820 | 1.789 | 2.366 | 2.457 | 1.443 | 0.026 | 0.027 | 0.017 | 0.014 | 1.000 |
| Maternal B Cells | 0.102 | 0.777 | -3.390 | -3.348 | 3.160 | 2.959 | 0.005 | 0.003 | 0.011 | 0.007 | 0.999 |
| Maternal Plasma Cells | 0.221 | 0.038 | 1.167 | 1.445 | 1.078 | 0.795 | 0.008 | 0.009 | 0.005 | 0.004 | 0.140 |
| Maternal Naive CD8+ T Cells | 0.164 | 0.222 | 2.083 | 2.871 | 2.620 | 1.558 | 0.040 | 0.049 | 0.028 | 0.030 | 0.479 |
| Fetal Extravillous Trophoblasts | 0.088 | 0.896 | 3.409 | 3.526 | 1.255 | 0.568 | 0.072 | 0.071 | 0.038 | 0.037 | 1.000 |

**Supplemental Table 5. PTD cell type composition.** The cell type composition of PTD compared to control placentas. The mean and standard deviation are reported for each group as clr-transformed and percentage cell composition. The p-values were calculated by Kolmogorov-Smirnov tests followed by Benjamini-Hochberg multiple hypothesis corrections.

| Cell Type | KS Statistic | p-value | Group Mean (CLR) | Control Mean (CLR) | Group STD (CLR) | Control STD (CLR) | Group Mean (%) | Control Mean (%) | Group STD (%) | Control STD (%) | p-value Adjusted |
|---|---|---|---|---|---|---|---|---|---|---|---|
| Fetal Mesenchymal Stem Cells | 0.252 | 0.068 | 3.556 | 3.693 | 0.340 | 0.286 | 0.067 | 0.076 | 0.024 | 0.022 | 0.209 |
| Fetal CD14+ Monocytes | 0.370 | 0.001 | -0.064 | 2.482 | 3.704 | 1.667 | 0.019 | 0.035 | 0.019 | 0.020 | 0.010 |
| Fetal CD8+ Activated T Cells | 0.412 | 0.000 | 3.626 | 3.262 | 0.297 | 0.914 | 0.071 | 0.054 | 0.019 | 0.020 | 0.002 |
| Fetal Naive CD4+ T Cells | 0.294 | 0.020 | -2.296 | -0.746 | 3.428 | 3.388 | 0.006 | 0.010 | 0.009 | 0.012 | 0.090 |
| Fetal Naive CD8+ T Cells | 0.191 | 0.278 | -4.367 | -5.196 | 2.270 | 0.000 | 0.002 | 0.000 | 0.011 | 0.000 | 0.527 |
| Fetal Natural Killer T Cells | 0.200 | 0.231 | -1.655 | -0.822 | 3.513 | 3.205 | 0.007 | 0.007 | 0.010 | 0.007 | 0.489 |
| Fetal B Cells | 0.497 | 0.000 | 2.990 | 3.364 | 0.510 | 0.307 | 0.040 | 0.054 | 0.014 | 0.014 | 0.000 |
| Fetal GZMK+ Natural Killer | 0.309 | 0.012 | -4.009 | -2.858 | 2.320 | 3.105 | 0.001 | 0.003 | 0.002 | 0.006 | 0.058 |
| Fetal Memory CD4+ T Cells | 0.273 | 0.038 | 3.655 | 3.828 | 0.356 | 0.248 | 0.074 | 0.085 | 0.022 | 0.018 | 0.140 |
| Fetal Hofbauer Cells | 0.412 | 0.000 | -0.523 | 1.500 | 3.037 | 1.816 | 0.008 | 0.015 | 0.014 | 0.011 | 0.002 |
| Fetal Plasmacytoid Dendritic Cells | 0.473 | 0.000 | 3.321 | 3.159 | 1.546 | 0.864 | 0.067 | 0.048 | 0.025 | 0.015 | 0.000 |
| Fetal GZMB+ Natural Killer | 0.309 | 0.012 | -5.196 | -5.130 | 0.000 | 0.690 | 0.000 | 0.000 | 0.000 | 0.001 | 0.058 |
| Fetal Endothelial Cells | 0.209 | 0.190 | 3.429 | 3.570 | 0.289 | 0.269 | 0.058 | 0.066 | 0.015 | 0.017 | 0.430 |
| Fetal Syncytiotrophoblast | 0.703 | 0.000 | -1.782 | 2.623 | 3.586 | 0.563 | 0.008 | 0.028 | 0.011 | 0.013 | 0.000 |
| Fetal Fibroblasts | 0.245 | 0.080 | -2.382 | -1.794 | 3.158 | 3.107 | 0.004 | 0.004 | 0.007 | 0.008 | 0.226 |
| Fetal Cytotrophoblasts | 0.524 | 0.000 | 5.063 | 5.229 | 0.266 | 0.097 | 0.294 | 0.339 | 0.061 | 0.032 | 0.000 |
| Fetal Proliferative Cytotrophoblasts | 0.309 | 0.012 | -5.196 | -5.196 | 0.000 | 0.000 | 0.000 | 0.000 | 0.000 | 0.000 | 0.058 |
| Fetal Nucleated Red Blood Cells | 0.455 | 0.000 | -1.993 | -3.827 | 2.175 | 1.868 | 0.001 | 0.000 | 0.001 | 0.000 | 0.000 |
| Maternal CD8+ Activated T Cells | 0.097 | 0.953 | -2.167 | -2.083 | 3.451 | 3.438 | 0.007 | 0.006 | 0.011 | 0.010 | 1.000 |
| Maternal Naive CD4+ T Cells | 0.227 | 0.125 | -2.553 | -3.920 | 3.368 | 2.673 | 0.006 | 0.003 | 0.017 | 0.010 | 0.322 |
| Maternal FCGR3A+ Monocytes | 0.309 | 0.012 | -5.196 | -5.196 | 0.000 | 0.000 | 0.000 | 0.000 | 0.000 | 0.000 | 0.058 |
| Maternal CD14+ Monocytes | 0.403 | 0.000 | 0.505 | -1.271 | 3.414 | 3.538 | 0.022 | 0.009 | 0.022 | 0.012 | 0.003 |
| Maternal Natural Killer Cells | 0.258 | 0.058 | 1.355 | 2.366 | 2.848 | 1.443 | 0.024 | 0.027 | 0.017 | 0.014 | 0.198 |
| Maternal B Cells | 0.582 | 0.000 | 0.714 | -3.348 | 3.205 | 2.959 | 0.020 | 0.003 | 0.017 | 0.007 | 0.000 |
| Maternal Plasma Cells | 0.330 | 0.006 | 1.171 | 1.445 | 0.560 | 0.795 | 0.007 | 0.009 | 0.004 | 0.004 | 0.034 |
| Maternal Naive CD8+ T Cells | 0.491 | 0.000 | -1.049 | 2.871 | 4.044 | 1.558 | 0.019 | 0.049 | 0.023 | 0.030 | 0.000 |
| Fetal Extravillous Trophoblasts | 0.597 | 0.000 | 4.435 | 3.526 | 0.490 | 0.568 | 0.171 | 0.071 | 0.077 | 0.037 | 0.000 |

**Supplemental Table 6. FGR+HDP cell type composition.** The cell type composition of FGR+HDP compared to control placentas. The mean and standard deviation are reported for each group as clr-transformed and percentage cell composition. The p-values were calculated by Kolmogorov-Smirnov tests followed by Benjamini-Hochberg multiple hypothesis corrections.

| Cell Type | KS Statistic | p-value | Group Mean (CLR) | Control Mean (CLR) | Group STD (CLR) | Control STD (CLR) | Group Mean (%) | Control Mean (%) | Group STD (%) | Control STD (%) | p-value Adjusted |
|---|---|---|---|---|---|---|---|---|---|---|---|
| Fetal Mesenchymal Stem Cells | 0.130 | 0.475 | 3.722 | 3.693 | 0.329 | 0.286 | 0.079 | 0.076 | 0.026 | 0.022 | 0.734 |
| Fetal CD14+ Monocytes | 0.280 | 0.003 | 1.719 | 2.482 | 2.550 | 1.667 | 0.025 | 0.035 | 0.016 | 0.020 | 0.023 |
| Fetal CD8+ Activated T Cells | 0.189 | 0.105 | 3.438 | 3.262 | 0.406 | 0.914 | 0.060 | 0.054 | 0.017 | 0.020 | 0.280 |
| Fetal Naive CD4+ T Cells | 0.143 | 0.354 | -1.321 | -0.746 | 3.500 | 3.388 | 0.008 | 0.010 | 0.011 | 0.012 | 0.616 |
| Fetal Naive CD8+ T Cells | 0.048 | 1.000 | -5.080 | -5.196 | 0.901 | 0.000 | 0.000 | 0.000 | 0.002 | 0.000 | 1.000 |
| Fetal Natural Killer T Cells | 0.100 | 0.777 | -0.797 | -0.822 | 3.282 | 3.205 | 0.008 | 0.007 | 0.008 | 0.007 | 0.999 |
| Fetal B Cells | 0.308 | 0.001 | 3.090 | 3.364 | 1.105 | 0.307 | 0.046 | 0.054 | 0.013 | 0.014 | 0.007 |
| Fetal GZMK+ Natural Killer | 0.158 | 0.248 | -3.567 | -2.858 | 2.781 | 3.105 | 0.002 | 0.003 | 0.005 | 0.006 | 0.499 |
| Fetal Memory CD4+ T Cells | 0.168 | 0.191 | 3.787 | 3.828 | 0.269 | 0.248 | 0.082 | 0.085 | 0.019 | 0.018 | 0.430 |
| Fetal Hofbauer Cells | 0.277 | 0.004 | 0.497 | 1.500 | 2.588 | 1.816 | 0.011 | 0.015 | 0.011 | 0.011 | 0.025 |
| Fetal Plasmacytoid Dendritic Cells | 0.188 | 0.106 | 3.217 | 3.159 | 1.139 | 0.864 | 0.054 | 0.048 | 0.020 | 0.015 | 0.280 |
| Fetal GZMB+ Natural Killer | 0.048 | 1.000 | -5.128 | -5.130 | 0.528 | 0.690 | 0.000 | 0.000 | 0.000 | 0.001 | 1.000 |
| Fetal Endothelial Cells | 0.078 | 0.952 | 3.581 | 3.570 | 0.281 | 0.269 | 0.067 | 0.066 | 0.017 | 0.017 | 1.000 |
| Fetal Syncytiotrophoblast | 0.330 | 0.000 | 1.832 | 2.623 | 2.024 | 0.563 | 0.022 | 0.028 | 0.015 | 0.013 | 0.003 |
| Fetal Fibroblasts | 0.139 | 0.390 | -2.249 | -1.794 | 3.291 | 3.107 | 0.005 | 0.004 | 0.009 | 0.008 | 0.648 |
| Fetal Cytotrophoblasts | 0.114 | 0.640 | 5.232 | 5.229 | 0.102 | 0.097 | 0.339 | 0.339 | 0.034 | 0.032 | 0.886 |
| Fetal Proliferative Cytotrophoblasts | 0.048 | 1.000 | -5.196 | -5.196 | 0.000 | 0.000 | 0.000 | 0.000 | 0.000 | 0.000 | 1.000 |
| Fetal Nucleated Red Blood Cells | 0.163 | 0.221 | -3.433 | -3.827 | 1.974 | 1.868 | 0.000 | 0.000 | 0.000 | 0.000 | 0.479 |
| Maternal CD8+ Activated T Cells | 0.112 | 0.661 | -2.526 | -2.083 | 3.373 | 3.438 | 0.005 | 0.006 | 0.009 | 0.010 | 0.890 |
| Maternal Naive CD4+ T Cells | 0.095 | 0.829 | -3.413 | -3.920 | 3.041 | 2.673 | 0.004 | 0.003 | 0.011 | 0.010 | 1.000 |
| Maternal FCGR3A+ Monocytes | 0.048 | 1.000 | -5.196 | -5.196 | 0.000 | 0.000 | 0.000 | 0.000 | 0.000 | 0.000 | 1.000 |
| Maternal CD14+ Monocytes | 0.274 | 0.004 | 0.502 | -1.271 | 2.952 | 3.538 | 0.014 | 0.009 | 0.013 | 0.012 | 0.025 |
| Maternal Natural Killer Cells | 0.092 | 0.854 | 2.282 | 2.366 | 1.617 | 1.443 | 0.027 | 0.027 | 0.014 | 0.014 | 1.000 |
| Maternal B Cells | 0.276 | 0.004 | -1.538 | -3.348 | 3.381 | 2.959 | 0.007 | 0.003 | 0.009 | 0.007 | 0.025 |
| Maternal Plasma Cells | 0.149 | 0.310 | 1.298 | 1.445 | 0.620 | 0.795 | 0.008 | 0.009 | 0.004 | 0.004 | 0.562 |
| Maternal Naive CD8+ T Cells | 0.231 | 0.025 | 2.221 | 2.871 | 2.363 | 1.558 | 0.038 | 0.049 | 0.027 | 0.030 | 0.103 |
| Fetal Extravillous Trophoblasts | 0.228 | 0.027 | 3.784 | 3.526 | 0.487 | 0.568 | 0.089 | 0.071 | 0.042 | 0.037 | 0.110 |

**Supplemental Table 7. PE cell type composition.** The cell type composition of PE compared to control placentas. The mean and standard deviation are reported for each group as clr-transformed and percentage cell composition. The p-values were calculated by Kolmogorov-Smirnov tests followed by Benjamini-Hochberg multiple hypothesis corrections.

| Feature | Variance Inflation Factor (VIF) Female Fetuses | Variance Inflation Factor (VIF) Male Fetuses |
|---|---|---|
| Gestational Weeks | 6.67 | 5.49 |
| Pregravid BMI$^2$ | 5.53 | 4.03 |
| Labor Initiation | 1.65 | 2.06 |
| Smoker | 2.19 | 2.03 |
| Illicit Drug User | 2.21 | 2.06 |

**Supplemental Table 8. Variance inflation factors for GLM features in female and male fetuses**.

**Supplemental Table 9. GLMs for female fetuses**. For each analyte the following model was fit: analyte ~ GestationalWeeks + PregravidBMI$^2$ + C(LaborInitiation) + C(Smoker) + C(IllicitDrugUser). Anderson-Darling tests were used to evaluate the distribution of the data. Z-score was used to detect outliers with the threshold set as the absolute value of three, if more than 2.5% of all measurements surpassed this threshold then the analyte was declared an outlier. If the analyte had a non-normal distribution and/or the analyte was an outlier than a Gamma family was used to train the GLMs, otherwise a Gaussian family was applied. Benjamini-Hochberg multiple hypothesis adjustment was performed. The β-coefficients for the variables, y-intercepts, p-value, adjusted p-value, and pseudo R-squared value for each GLM were reported. This table takes up more than one page, so it is provided as an .xlsx file available for download.

**Supplemental Table 10. GLMs for male fetuses**. For each analyte the following model was fit: analyte ~ GestationalWeeks + PregravidBMI$^2$ + C(LaborInitiation) + C(Smoker) + C(IllicitDrugUser). Anderson-Darling tests were used to evaluate the distribution of the data. Z-score was used to detect outliers with the threshold set as the absolute value of three, if more than 2.5% of all measurements surpassed this threshold then the analyte was declared an outlier. If the analyte had a non-normal distribution and/or the analyte was an outlier than a Gamma family was used to train the GLMs, otherwise a Gaussian family was applied. Benjamini-Hochberg multiple hypothesis adjustment was performed. The β-coefficients for the variables, y-intercepts, p-value, adjusted p-value, and pseudo R-squared value for each GLM were reported. This table takes up more than one page, so it is provided as an .xlsx file available for download.

| | Control n=30 | FGR n=30 | FGR+HDP n=30 | PE n=30 | PTD n=30 |
|---|---|---|---|---|---|
| Edges (n) | 1 | 10 | 80,612 | 4,032 | 3,020 |
| Nodes (n) | 2 | 17 | 714 | 1625 | 1161 |

**Supplemental Table 11. Network and partition quality metrics and overview after downsampling.** Random downsampling was performed to select thirty placentas for each condition. Then interomics correlations followed by Bonferroni correction as described. The number of edges and nodes participating in significant interomics correlations are reported by condition.

**Supplemental Table 12. Control interomics communities**. For each node of the control interomics network we report the community to which it belongs along with its closeness centrality score, which was normalized to a scale of 0 to 1 for each community. This table takes up more than one page, so it is provided as an .xlsx file available for download.

**Supplemental Table 13. FGR interomics communities**. For each node of the FGR interomics network we report the community to which it belongs along with its closeness centrality score, which was normalized to a scale of 0 to 1 for each community. This table takes up more than one page, so it is provided as an .xlsx file available for download.

**Supplemental Table 14. PTD interomics communities**. For each node of the PTD interomics network we report the community to which it belongs along with its closeness centrality score, which was normalized to a scale of 0 to 1 for each community. This table takes up more than one page, so it is provided as an .xlsx file available for download.

**Supplemental Table 15. FGR+HDP interomics communities**. For each node of the FGR+HDP interomics network we report the community to which it belongs along with its closeness centrality score, which was normalized to a scale of 0 to 1 for each community. This table takes up more than one page, so it is provided as an .xlsx file available for download.

**Supplemental Table 16. PE interomics communities**. For each node of the PE interomics network we report the community to which it belongs along with its closeness centrality score, which was normalized to a scale of 0 to 1 for each community. This table takes up more than one page, so it is provided as an .xlsx file available for download.

| Library or Language | Version | Citation | Documentation |
| --- | --- | --- | --- |
| Jupyter | 6.5.4 | Kluyver et al. 2016[2] | https://jupyter.org/ |
| logging | 0.5.1.2 | open source | https://docs.python.org/3/library/logging.html |
| matplotlib | 3.8.2 | Hunter et al. 2007[3] | https://matplotlib.org/ |
| networkx | 3.1 | Hagberg et al. 2008[4] | https://networkx.org/ |
| nxviz | 0.7.3 | open source | https://pypi.org/project/nxviz/ |
| numpy | 1.24.3 | Harris et al. 2020[5] | https://numpy.org/ |
| pandas | 2.0.1 | Reback et al. 2020[6] | https://pandas.pydata.org/ |
| python | 3.11.6 | open source | https://docs.python.org/release/3.11.6/ |
| pyvis | 0.3.1 | open source | https://pyvis.readthedocs.io/en/latest/documentation.html |
| scipy | 1.10.1 | Virtanen et al. 2020[7] | https://scipy.org/ |
| seaborn | 0.12.2 | Waskom et al. 2021[8] | https://seaborn.pydata.org/index.html |
| sklearn | 1.2.2 | Pedregosa et al. 2011[9] | https://scikit-learn.org/stable/index.html |
| statsmodels | 0.14.0 | Seabold & Perktold 2010[10] | https://www.statsmodels.org/stable/index.html |

**Supplemental Table 17. Key resources: software and algorithms.**
